# Open science resources for the discovery and analysis of *Tara* Oceans Data

**DOI:** 10.1101/019117

**Authors:** Stéphane Pesant, Fabrice Not, Marc Picheral, Stefanie Kandels-Lewis, Noan Le Bescot, Gabriel Gorsky, Daniele Iudicone, Eric Karsenti, Sabrina Speich, Romain Troublé, Céline Dimier, Sarah Searson, *Tara* Oceans Consortium Coordinators

## Abstract

The *Tara* Oceans expedition (2009-2013) sampled contrasting ecosystems of the world oceans, collecting environmental data and plankton, from viruses to metazoans, for later analysis using modern sequencing and state-of-the-art imaging technologies. It surveyed 210 ecosystems in 20 biogeographic provinces, collecting over 35000 samples of seawater and plankton. The interpretation of such an extensive collection of samples in their ecological context requires means to explore, assess and access raw and validated data sets. To address this challenge, the Tara Oceans Consortium offers open science resources, including the use of open access archives for nucleotides (ENA) and for environmental, biogeochemical, taxonomic and morphological data (PANGAEA), and the development of on line discovery tools and collaborative annotation tools for sequences and images. Here, we present an overview of Tara Oceans Data, and we provide detailed registries (data sets) of all campaigns (from port-to-port), stations and sampling events.

## Background & Summary

Over many centuries, global expeditions have led to major scientific breakthroughs, notably with the early voyages of the H.M.S. Beagle (1831-1836) and the H.M.S. Challenger (1872-1876). Ocean exploration now provides promising first steps towards understanding the role of the ocean in global biogeochemical cycles and the impact of global climate change on ocean processes and marine biodiversity. Recently, the *Sorcerer II* expeditions (2003-2010) (Ref. 1) and the Malaspina expedition (2010-2011) (Ref. 2) carried out global surveys of prokaryotic metagenomes from the ocean’s surface and bathypelagic layer (>1000 m), respectively. The *Tara* Oceans Expedition (2009-2013) complemented these surveys by collecting a wide variety of planktonic organisms (from viruses to fish larvae) from the ocean’s surface (0-200 m) and mesopelagic zone (200-1000m) at a global scale. Overall, *Tara* Oceans surveyed 210 ecosystems in 20 biogeographic provinces, collecting over 35000 samples of seawater and plankton. Organising such a knowledge base is essential to safeguard, discover and share *Tara* Oceans data. To address this challenge, *Tara* Oceans offers open science resources, including the use of open access data archives and the development of online tools for the collaborative annotation of sequences and images, and the discovery of *Tara* Oceans data.

*Tara* Oceans adopts the principle of open access and early release of raw and validated data sets. In the case of molecular data, raw short sequence reads are archived at the European Bioinformatics Institute *short read archive* (<u>http://www.ebi.ac.uk/ena/</u>) and made available immediately after manual curation of metadata. More advanced data (assemblies, annotations, etc.) will be released immediately after validation and before publication, and other versions will be released when available. In the case of environmental, biogeochemical, taxonomic and morphological measurements, data are published at PANGAEA, Data Publisher for Earth and Environmental Science (<u>http://www.pangaea.de</u>) and made available immediately after manual curation of metadata.

By combining modern sequencing and state-of-the-art imaging technologies, *Tara* Oceans is at the cutting edge of marine science (3). The amount of data generated by these technologies is unprecedented in the field of plankton ecology and requires adapted storage infrastructures and collaborative platforms to carry out manual and automated annotation of sequences and high throughput images. These open science resources are currently being developed by *Tara* Oceans.

A first series of publications has demonstrated the potential of *Tara* Oceans data to study the ecology of plankton and the structural and functional diversity of viruses, prokaryotes and eukaryotes in the global ocean (4-11). These publications are based on a fraction of the samples analysed so far and thus represent only the tip of the iceberg. The exploration of Tara Oceans data by the scientific community will undoubtedly lead to new hypotheses and emerging concepts in domains unforeseen by the Tara Oceans Consortium. The current discovery portal of *Tara* Oceans offers a simple map interface that links each sampling location to available environmental and molecular data (<u>http://www.taraoceans-dataportal.org/</u>). It will however evolve to offer advanced search functionalities based on geospatial, methodological, environmental, morphological, taxonomic, phylogenetic and ecological criteria.

Here, we present an overview of the sampling strategy and size-fractionation approach of the *Tara* Oceans Expedition (***Methods Section***) and we explain the rationale behind the choice of sampling devices (***Technical Validation Section***). Most importantly, we provide registries (data sets) describing all campaigns (from port-to-port), stations and sampling events (***Data Records Section***). These registries contain geospatial, temporal and methodological information that will be essential for researchers to explore and assess the quality of *Tara* Oceans data. Environmental data sets are already available openly, in whole or in part, and additional data sets will be progressively released to the community. We intend to submit additional publications describing specific data types (e.g.,Data Citations 1-5) in more detail, further extending the value of this resource as the data becomes available.

## Methods

As a research infrastructure, the *Tara* Oceans Expedition mobilised over 100 scientists to sample the world oceans on board a 36 m long schooner (SV *Tara*) refitted to operate state-of-the-art oceanographic equipment (Figure 1). On board the schooner, the team was consistently composed of five sailors and six scientists, including one chief scientist, two oceanography engineers in charge of deck operations, instrument maintenance and data management, two biology engineers preparing and preserving samples for later morphological and genetic analyses, and one optics engineer in charge of imaging live samples on board. A winch equipped with 2400 m of cable was installed to deploy sampling devices from the stern of the ship, and an industrial peristaltic pump was installed on starboard to sample large volumes of water from various depths down to 60 m. Peristaltic and vacuum filtration systems used to concentrate plankton on membranes of various pore sizes were setup in a laboratory container (wet lab) located outside on port side. Flow-through instruments connected to the continuous surface sampling system were installed in the fore peak and in a laboratory (dry lab) inside the schooner at the centre of the ship on port side.

**Figure 1.**
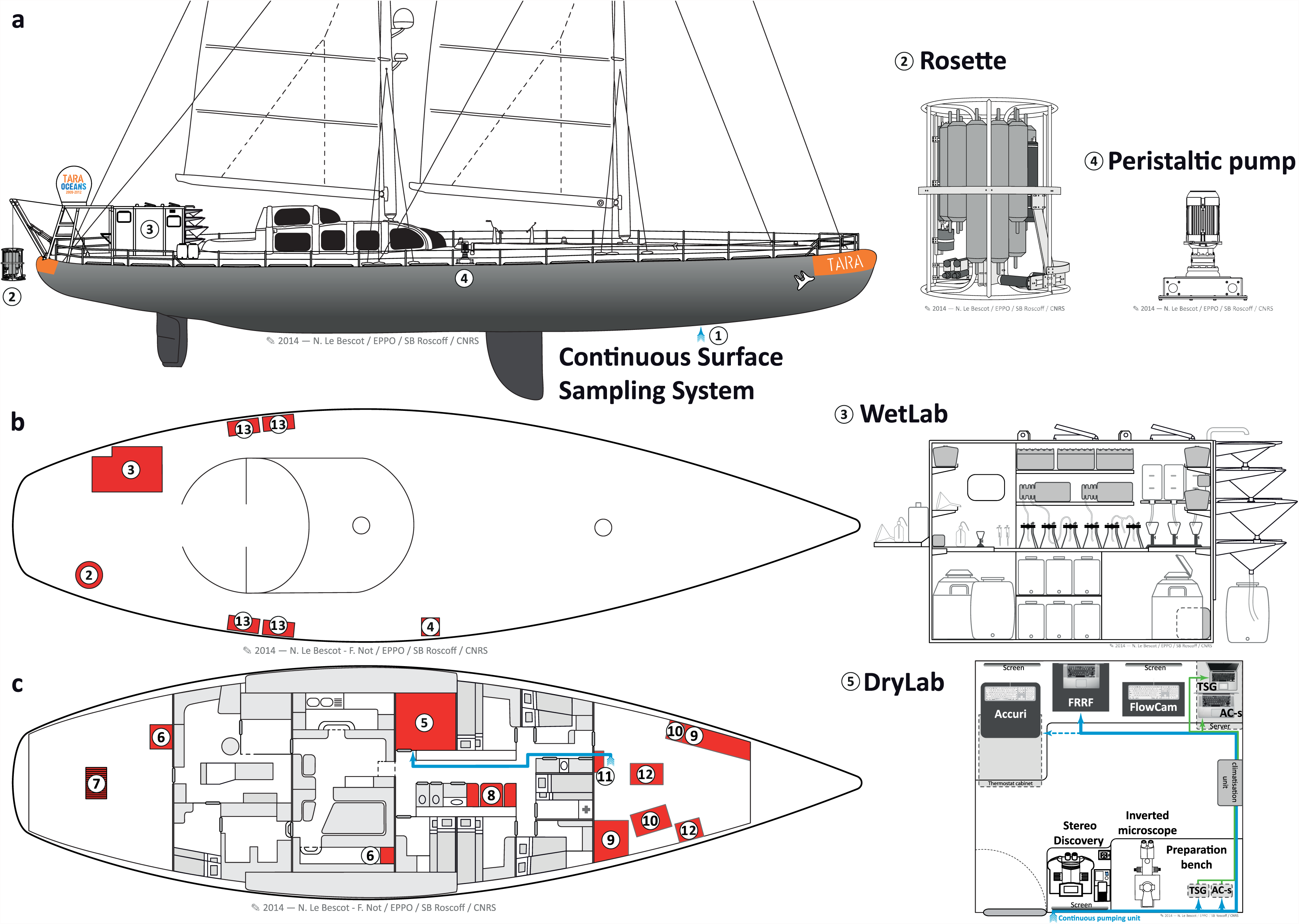
Sampling devices and working areas on-board SV *Tara.* Sampling devices and working areas on-board SV *Tara* are shown from the vessel’s **[a]** side-view, **[b]** bird’s-eye-view of the deck, and **[c]** inside-view. They consist of the, **[1]** Continuous Surface Sampling System [CSSS]; **[2]** Rosette Vertical Sampling System [RVSS]; **[3]** wet lab; **[4]** High Volume Peristaltic pump [HVP-PUMP]; **[5]** dry lab; **[6]** oceanography engineers data acquisition and processing area; **[7]** winch; **[8]** video imaging area; **[9]** storage areas at room temperature; **[10]** storage areas at +4°C and −20°C; **[11]** MilliQ water system and AC-s system; **[12]** diving equipment, flowcytobot and ALPHA instruments; and **[13]** storage boxes. The flow of seawater from the continuous surface sampling system to the dry lab is shown in blue.

The sampling strategy and methodology of the *Tara* Oceans Expedition is presented in six sub-sections. The first four describe why and how the environmental context was determined **[1]** at the mesoscale using remote sensing and meteorological data; **[2]** from sensors mounted on the continuous surface water sampling system; **[3]** from sensors mounted on the vertical profile sampling system; and **[4]** from discrete water samplers (Niskin bottles) mounted on the vertical profile sampling system. The last two Sub-Sections describe how **[5]** environmental features were selected and sampled; and how **[6]** plankton were collected for imaging and genetic analyses. These methods were also described briefly in (3).

### [1] Atmospheric and oceanographic context at the mesoscale

The regular sampling programme was designed to study a variety of marine ecosystems and to target well-defined meso- to large-scale features such as gyres, eddies, currents, frontal zones, upwellings, hot spots of biodiversity, low pH or low oxygen concentrations. A total of 210 stations were characterised at the mesoscale to provide richer environmental context for the morphological and genomic study of plankton (Figure 2). In order to identify these features before sampling but also to assess *a posteriori* if sampling events carried out during a station were taken within a relatively homogeneous environment, the atmospheric and oceanographic context were determined at the mesoscale, using climatologies, remote sensing products and arrays of Argo profiling floats. Meteorological forecast services, satellite observations (Chlorophyll *a*, sea surface temperature (SST) and altimetry) and real-time ocean model outputs (Mercator Ocean) were also used on a daily basis to revise sampling positions with respect to the selected oceanographic features.

**Figure 2.**
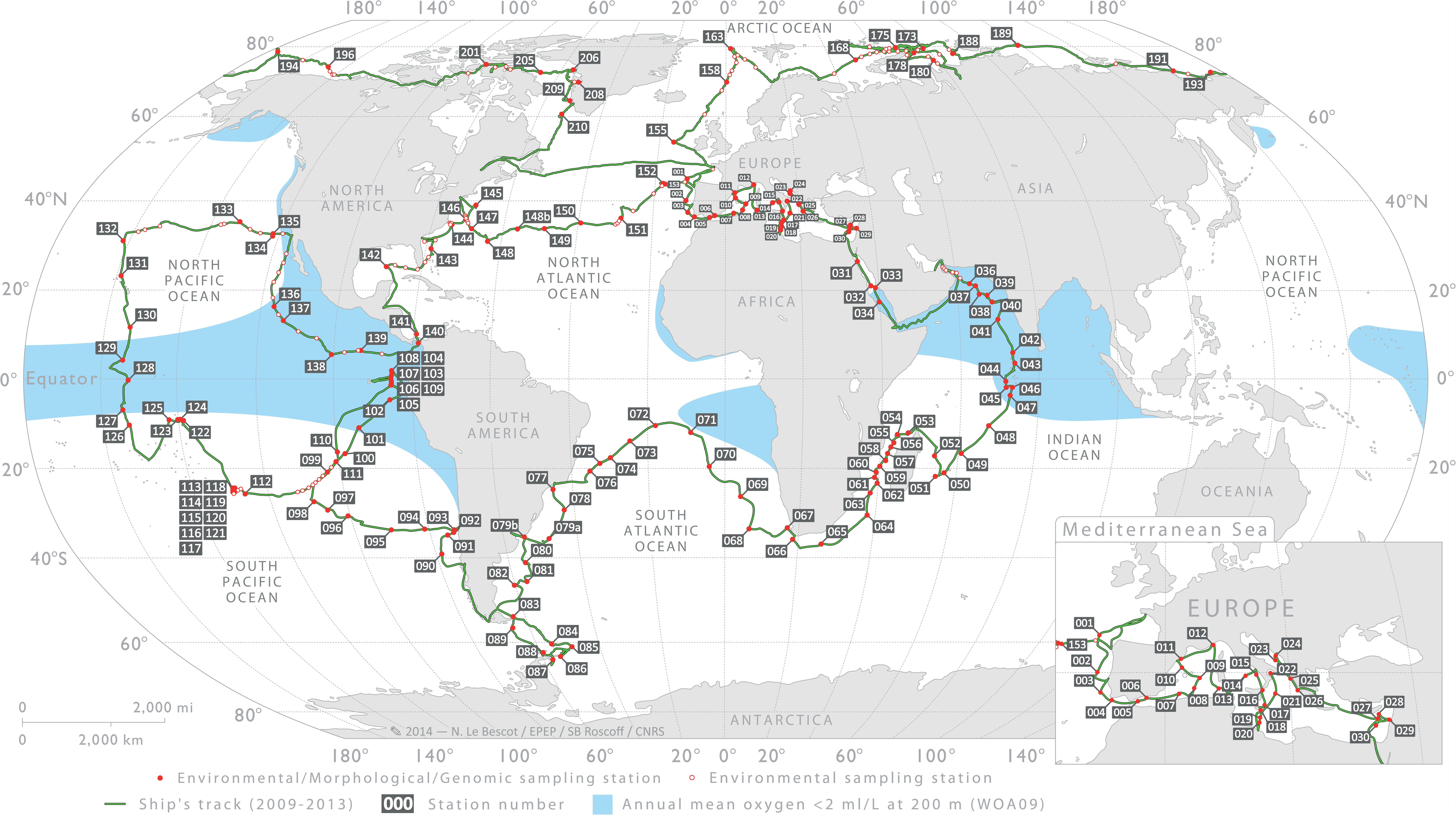
Sampling route and stations of the *Tara* Oceans Expedition. Sampling route of the *Tara* Oceans Expedition (green track), showing station labels and areas (blue shade) where the annual mean oxygen concentration is <2 ml/L (WOA09), usually corresponding also to high CO_2_ concentration and low pH. Detailed information about each station is given in (Data Citation 7).

Mapped altimetry from AVISO (Archiving Validation and Interpretation of Satellite Data in Oceanography), mapped operational SST (OSTIA), and satellite ocean colour (ACRI-ST GlobColour service) were used to describe the spatial and temporal variability of key environmental parameters at each sampling station. In addition, Temperature-Salinity profiles available around sampling stations were compiled from the Argo autonomous network array. Finally, a [BATOS] meteorological station mounted on-board *Tara* continuously measured wind speed and direction, and air temperature, pressure and humidity, which helped determine the variability of atmospheric conditions and vertical mixing of surface waters.

In addition to the regular sampling programme, topical experiments were designed to study ocean processes that operate at spatial and/or temporal scales larger or smaller than the mesoscale (Figure 3). Topics included, for example, diurnal processes, storm-induced perturbation of community structure and functions, latitudinal diversity gradients, oxygen minimum zone (6), island effects on iron fertilization, and longitudinal transport by Agulhas rings across the South Atlantic Ocean (11). For topical experiments, oceanographic context was sometimes enriched by using automated underwater vehicles (e.g., gliders (12), ProvBio (13)), surface-tethered Argo drifters, lowered ADCP mounted on the rosette, basin scale eddy-field simulations and climatologies, and state-of-the-art physical models of global ocean circulation with biogeochemistry and genome-informed models of microbial processes (14). The specific sampling strategy of each topical experiment is available in the respective cruise summary reports (see ***Data Records Section***). *Tara* Oceans data corresponding to methods described in this section are in part already open to the public at PANGAEA (Data Citation 1).

**Figure 3.**
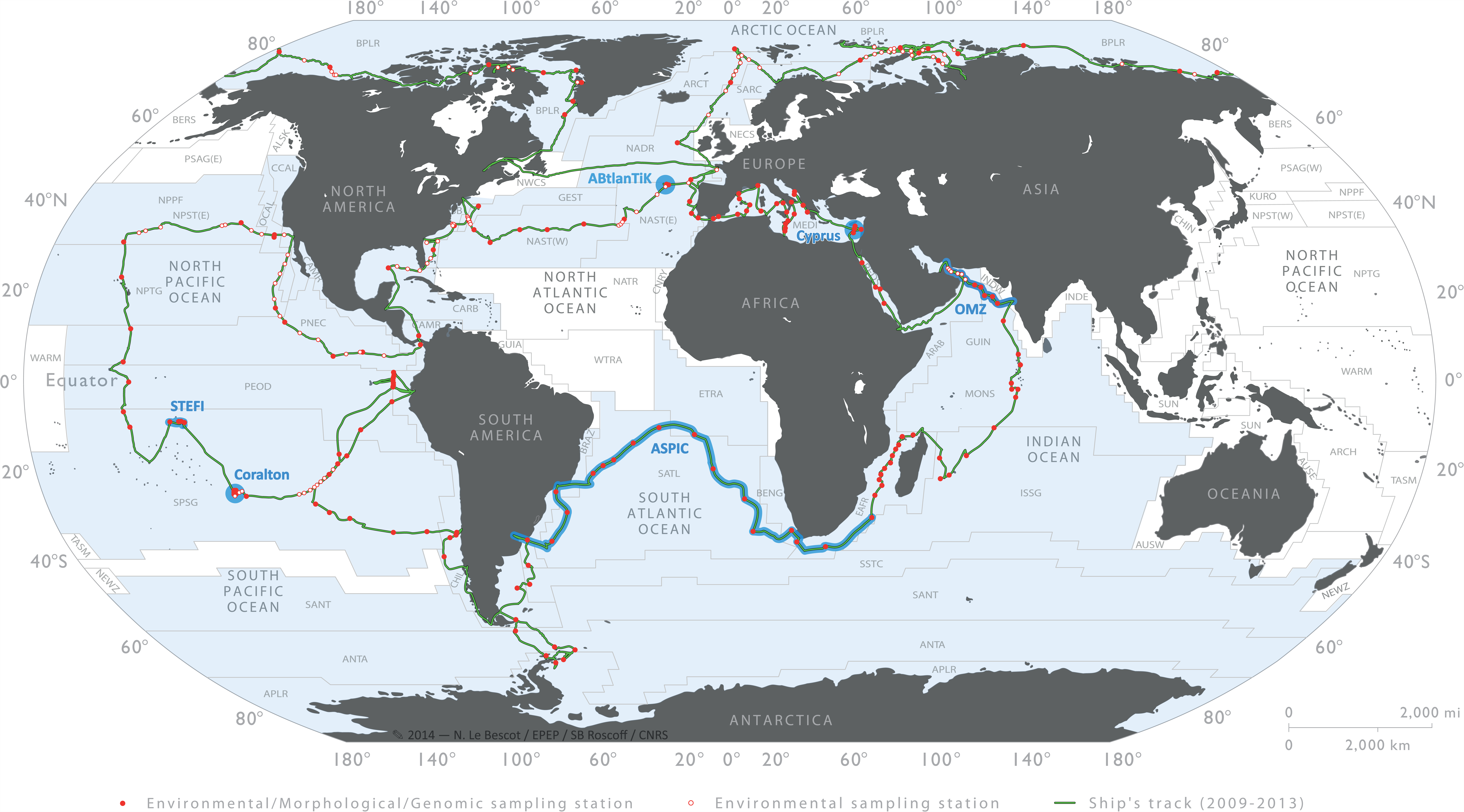
Sampling route, stations and topical experiments of the *Tara* Oceans Expedition. Sampling route of the *Tara* Oceans Expedition (green track), showing stations where plankton were sampled in their environmental context (full red dots) and where only environmental conditions were measured (open red dots). Topical experiments are identified along the sampling route (light blue). Longhurst biogeographical provinces (34) are shown in the background and those sampled during *Tara* Oceans Expedition are highlighted in blue.

### [2] Properties of seawater and particulate matter from physical, optical and imaging sensors mounted on the continuous surface water sampling system

Continuous measurements of surface water physical, chemical and biological properties serve the dual purpose of a) assessing the boundaries and the homogeneity/heterogeneity of an ecosystem studied during a station, and b) assessing the connectivity between stations. Underway measurements were often used to fine tune the location of sampling stations that were initially selected based on satellite images.

The in-line, Continuous Surface Sampling System [CSSS] installed on SV *Tara* (15,000 miles long track) comprised a SeaBird [TSG] temperature and conductivity sensor, a WETLabs [AC-S] spectrophotometer, a WETLabs chlorophyll [Fluorometer], and a Fast Repetition Rate Fluorometer [FRRF] to assess photosynthetic efficiency. All data were recorded simultaneously and archived daily in a single file, including navigation, date/time and GPS position. Water was pumped at the front of the vessel from ˜2m depth, then de-bubbled and circulated through the [AC-S], [TSG], [Fluorometer], and [FRRF]. An automated switching system provided periodic 0.2μm filtered samples to the [AC-S], such that its particulate optical properties were not affected by instrument drift (15). Systems maintenance (instrument cleaning, flushing) was done approximately once a week and in port between successive campaigns. In the Arctic Ocean and Arctic Seas (2013 campaigns), additional sensors for pH, PCO_2_, optical backscattering (3 wavelengths), fluorescence emission [ALFA] and surface Photosynthetically Active Radiation [PAR] were added to the in-line system. A [FlowCytobot] also recorded images of microplankton every 20 minutes. Using daily discrete measurements of CDOM absorption with an [UltraPath] system, we calibrated the [AC-S] to also provide hourly CDOM absorption (besides particulate absorption and attenuation). Data were processed, quality-controlled, and are consistent with remote sensing. A total of 60 tracks of continuous measurements corresponding to methods described in this section are in large part already open to the public at PANGAEA (Data Citation 2-3).

### [3] Properties of seawater and particulate & dissolved matter from physical, optical and imaging sensors mounted on the vertical profile sampling system

Repeated deployments of a Rosette Vertical Sampling System [RVSS] during day and night also served the dual purpose of a) assessing the boundaries and the homogeneity/heterogeneity of mesoscale features during a station, and b) assessing the connectivity between stations. These deployments were essential to locate features that have a vertical component and have a signature below the surface, such as eddies, upwellings, fronts, deep chlorophyll maxima, and ***oxygen minimum zone***s. The [RVSS] was specifically designed with various sensors, comprising 2 pairs of conductivity and temperature sensors (Sea-Bird), chrorophyll and CDOM fluorometers (WETLabs), a 25 cm transmissiometer for particles 0.5-20 μm (WETLabs), a one-wavelength backscatter meter for particles 0.5-20 μm (WETLabs), and a Underwater Vision Profiler (16) for particles >100 μm and zooplankton >600 μm (Hydroptic). A sbe43 oxygen sensor (Sea-Bird) and an In Situ Ultraviolet Spectrophotometer (ISUS) nitrate sensor (SATLANTIC) were also mounted on the Rosette. In the Arctic Ocean and Arctic Seas (2013 campaigns), a second sbe43 oxygen sensor (Sea-Bird) and a four frequency acoustic profiler (Aquascat) were added. Each component was powered on specific Li-Ion batteries and CTD data were self-recorded at 24Hz. All sensors were calibrated in factory before, during and after the four year programme. Oxygen data were validated using climatologies. Nitrate and Fluorescence data were adjusted with discrete measurements from Niskin bottles mounted on the Rosette, and dark calibrations of the optical sensors were performed monthly on-board. A total of 837 vertical profiles were made during the Expedition. Additional stand-alone Sea-Bird components [sbe19] and [sbe9S] were exceptionally mounted directly on the oceanographic cable during harsh sea conditions, when the deployment of the rosette was not safe. In addition, apparent optical properties of sea water were measured using a surface tethered [RADIOMETRE-TSRB] in 2009-2012 and a profiling [RADIOMETRE-COPS] in 2013. Finally, a total of 101 deployments of the Secchi disk [SECCHI] provide a valuable, world-wide contribution to historical records of this very simple and yet fundamental sampling device. *Tara* Oceans data corresponding to methods described in this section are already open to the public at PANGAEA (Data Citation 4).

### [4] Properties of seawater and particulate & dissolved matter from discrete water samples

In addition to sensors mounted on the Rosette Vertical Sampling System [RVSS], seawater was collected using Niskin bottles [NISKIN] (6 × 8-L Niskins and 4 × 12-L Niskins) in order to further characterise environmental conditions in the ecosystem under study. Measurements include pigment concentrations from HPLC analysis (10 depths per vertical profile; 25 pigments per depth), the carbonate system (Surface and 400m; pHT, CO_2_, pCO_2_, fCO_2_, HCO_3_, CO_3_, Total alkalinity, Total carbon, OmegaAragonite, OmegaCalcite, and quality Flag), nutrients (10 depths per vertical profile; NO_2_, PO_4_, N0_2_/NO_3_, SI, quality Flags), DOC, CDOM, and dissolved oxygen isotopes. More than 200 vertical profiles of these properties were made across the world ocean. DOC, CDOM and dissolved oxygen isotopes are available only for the Arctic Ocean and Arctic Seas (2013 campaigns). *Tara* Oceans data corresponding to methods described in this section are already open to the public at PANGAEA (Data Citation 5).

### [5] Environmental features and sampling stations

During the *Tara* Oceans Expedition (2009-2013), plankton were sampled from 5-10-m thick layers in the water column, corresponding to specific environmental features that were characterised on-board from sensor measurements. Environmental features are defined by controlled vocabularies in the environmental ontology (EnvO; <u>http://environmentontology.org/</u>) (17).

The ***surface water layer*** (ENVO:00002042), sometimes labelled in the literature and databases as “surface”, “SRF”, “SUR”, “SURF” or “S”, was simply defined as a layer between 3 and 7 m below the sea surface. The ***deep chlorophyll maximum layer*** (ENVO:01000326), often labelled in the literature and databases as “DCM” or “D”, was determined from the chlorophyll fluorometer (WETLabs optical sensors) mounted on the Rosette Vertical Sampling System [RVSS]. The presence of a DCM may indicate a maximum in the abundance of plankton bearing chlorophyll pigments, or it may result from the higher chlorophyll content of plankton living in a darker environment (18). This can be assessed *a posteriori* using water samples analysed for pigments by HPLC methods and from plankton counts. The ***mesopelagic zone*** (ENVO:00000213), also labelled in the literature and databases as “MESO” or “M”, corresponds to the layer between 200 and 1000 m depths. The sampling depth within the ***mesopelagic zone*** was selected based on vertical profiles of temperature, salinity, fluorescence, nutrients, oxygen, and particulate matter. The selected depth varied from station to station, targeting for example a nutricline, a minimum concentration of oxygen, a maximum concentration of particulate matter, or a fixed depth of ca. 400 m when no particular feature could be identified. Other environmental features of special scientific interest include the ***oxygen minimum zone*** (ENVO:01000065), often labelled in the literature and data sets as “OMZ” or “O”, and the ***epipelagic mixing layer*** (ENVO:01000061), also labelled in the literature and data sets as “ML”, “MIX” or “X”.

A complete sampling station consisted of collecting plankton from three distinct environmental features, typically the ***surface water layer***, ***deep chlorophyll maximum layer***, and ***mesopelagic zone*** (Figure 4). Such a station lasted typically 24-48 hours and special care was taken to reposition *SV Tara* in order to remain within a radius of 10 km and sample a homogeneous ecosystem as much as possible (see previous two sub-sections). The sequence of sampling deployments varied but generally followed the order illustrated in Figure 4, i.e. ***surface water layer*** and ***mesopelagic zone*** during daytime on the first day, night sampling over fixed depths, and ***deep chlorophyll maximum layer*** during daytime on the second day. Sampling devices consisted essentially of a High Volume Peristaltic pump [HVP-PUMP], a Rosette Vertical Sampling System [RVSS] equipped with sensors and [NISKIN] bottles, instrumented plankton nets [NET*-TYPE-MESH*], a [SECCHI] disc, and a [RADIOMETRE] (Table 1).

**Figure 4.**
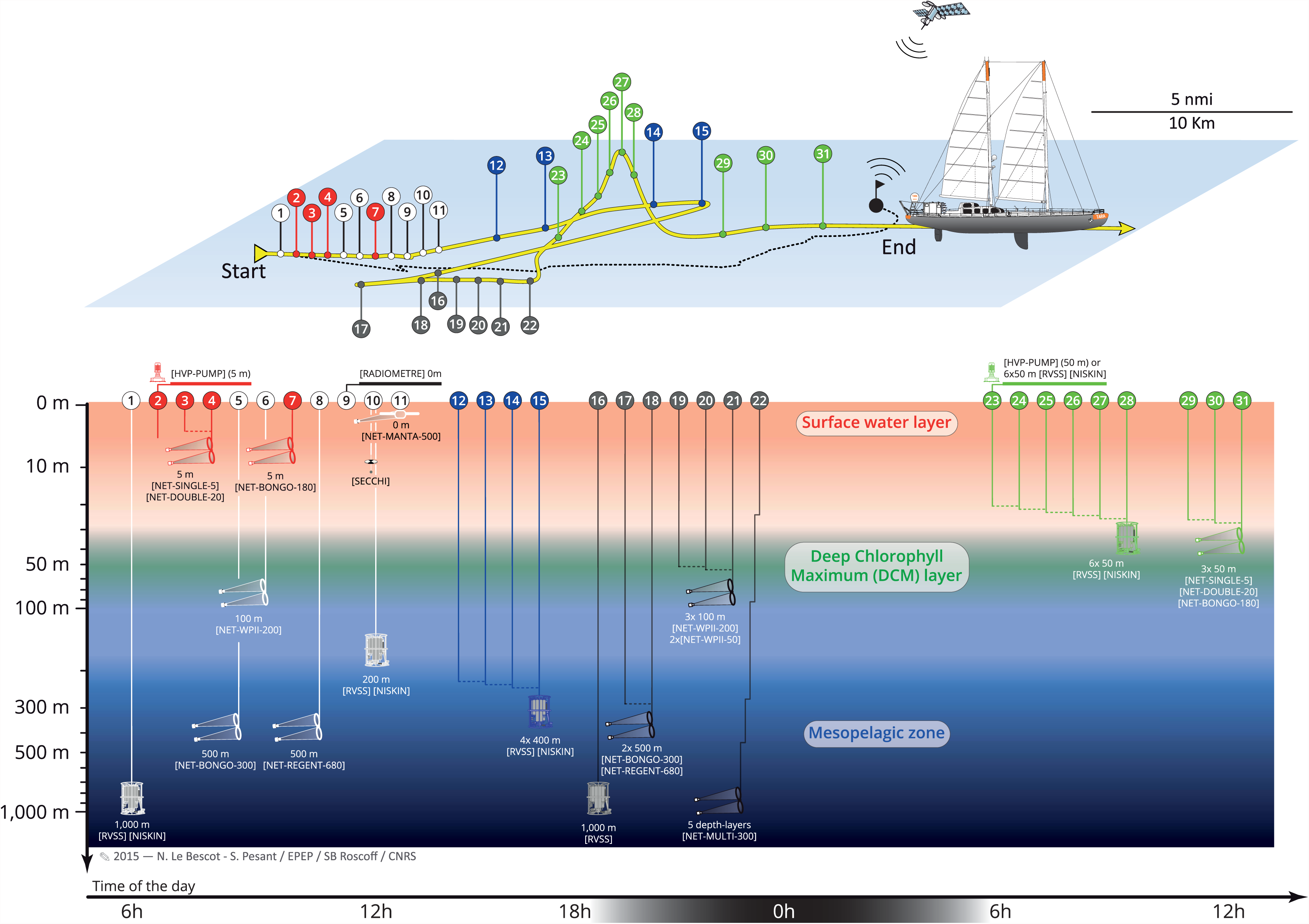
Spatial representation and chronology of sampling events during a 48 hours station. Coloured markers along the route of *SV Tara* (yellow surface track) correspond to sampling events targeting the ***surface water layer*** (red,), ***deep chlorophyll maximum layer*** (green, here at 50 m), and the ***mesopelagic zone*** (blue, here at 400 m). At some stations, an Argo drifter (10-m floating anchor and satellite positioning) was used to follow the water mass during sampling (black surface track). White and grey markers correspond to day and night time deployments, respectively, of plankton nets [*TYPE-MESH*] and rosette [RVSS] casts that covered fixed depth layers of 0-100m, 0-500m or 0-1000m.

**Table 1.**
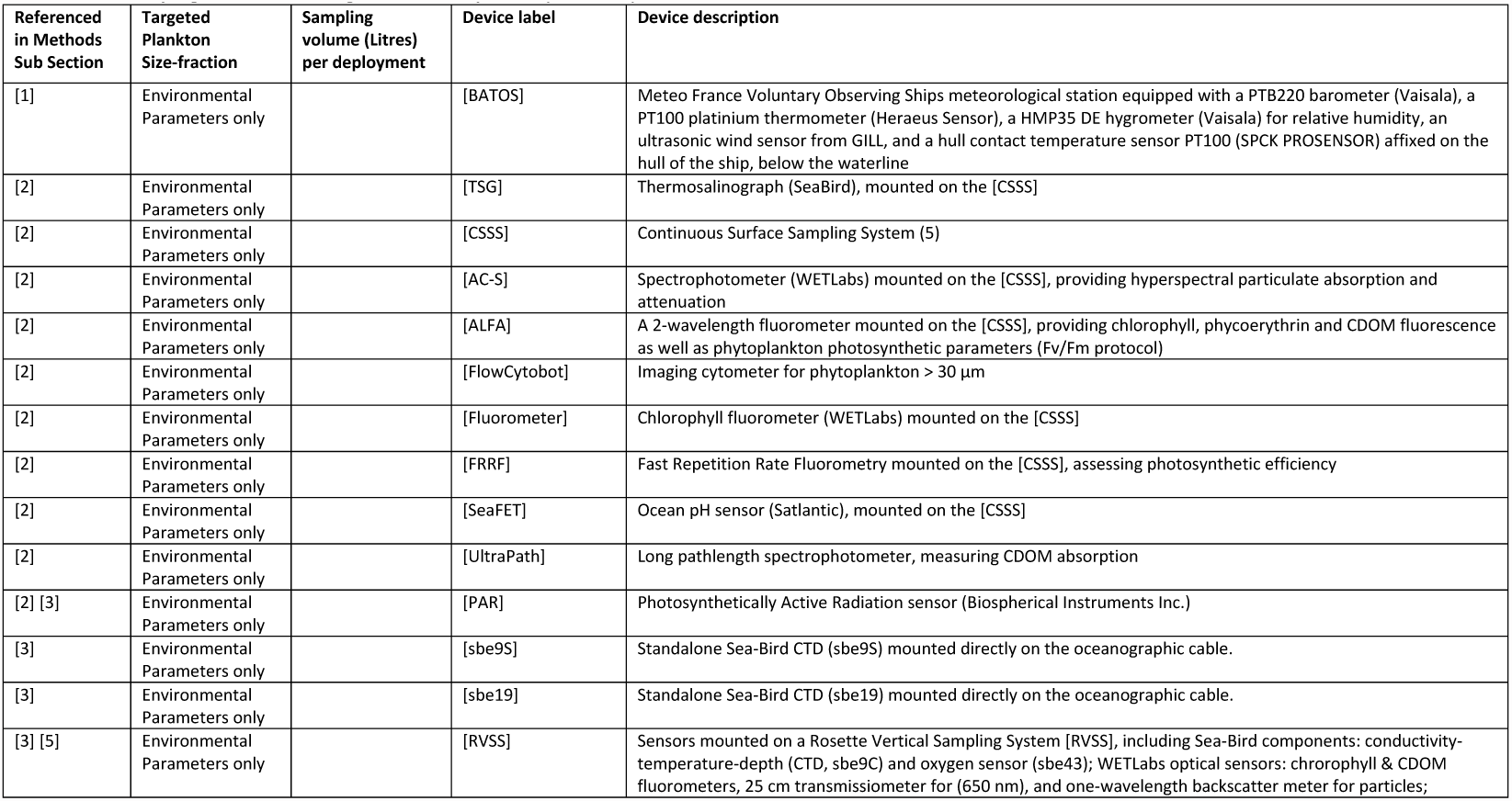
List of sampling devices used during *Tara* Oceans Expedition (2009-2013)

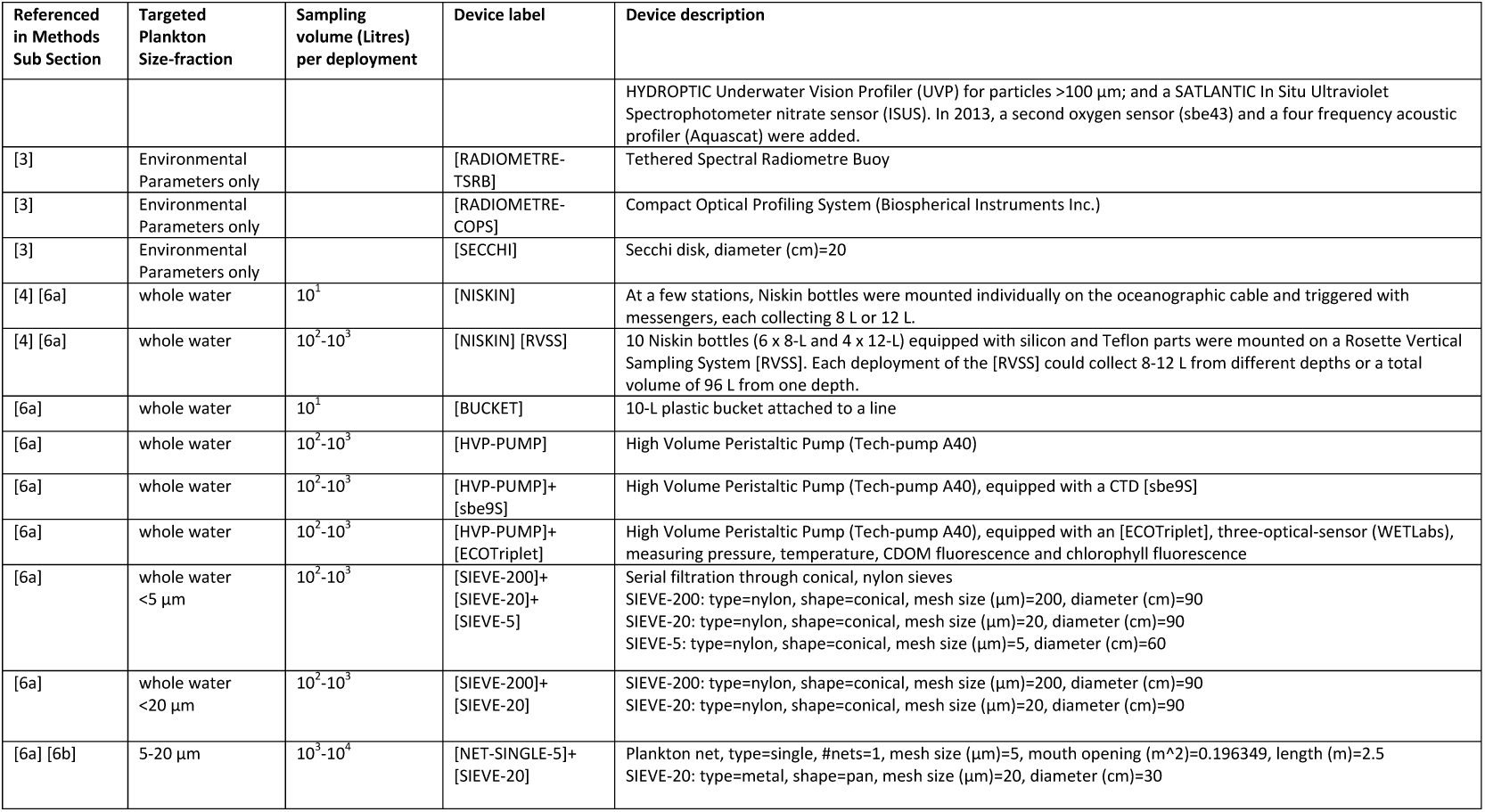

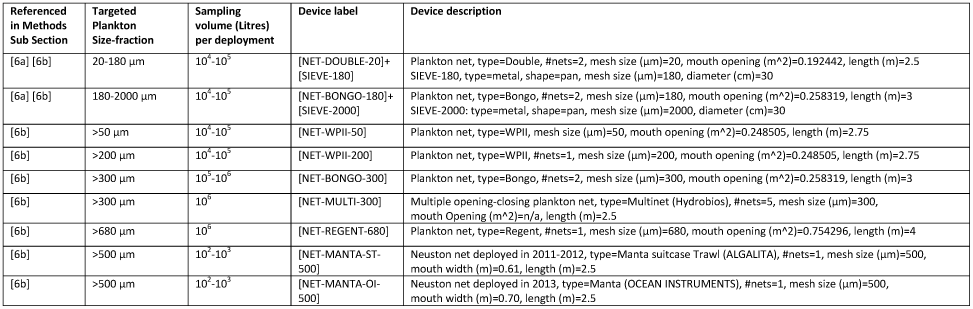

Plankton were sampled from a total of 210 stations, of which: 51 stations did not target a specific environmental feature and conducted classical vertical profiles of physical and optical sensors and depth integrated net tows; 57 stations sampled only the ***surface water layer***; 62 stations sampled the ***surface water layer*** and a second depth-specific feature; and 40 stations sampled the ***surface water layer***, the ***deep chlorophyll maximum layer*** and a third depth-specific feature. *Tara* Oceans data corresponding to methods described in this section are already open to the public at PANGAEA (Data Citations 6-8) and are described in the ***Data Records Section*** of the present paper.

### [6] Marine plankton

Plankton sampled during the *Tara* Oceans Expedition cover six orders of magnitude in size (10^-2^-10^5^ μm) and correspond to viruses, giant viruses (giruses), prokaryotes (bacteria and archaea), unicellular eukaryotes (protists), and multicellular eukaryotes (metazoans). These five groups form the bulk of biomass throughout the oceans and drive the global biogeochemical cycles that regulate the Earth system (19) (20) (21). Ocean viruses play an important role in plankton ecology by inducing mortality, horizontal gene transfer, and modulating microbial metabolism (22). They are thought to target diverse prokaryotic and eukaryotic hosts including microalgae and heterotrophic protists, and to play a role in the evolution of their hosts (23). Small viruses (<0.2 μm) are known to be ubiquitous and the most abundant plankton in seawater, while the larger *giant viruses* or *giruses* (0.2-1 μm) were discovered more recently and are increasingly observed in marine samples (24) (25). Prokaryotes are believed to be responsible for 30% of primary production and 95% of community respiration in oceans (26) and are thus a fundamental component of marine food webs and biogeochemical processes. They are often divided and studied as two size fractions: the free-living prokaryotes range in size from 0.22-3 μm, and those that are attached to larger cells, particles, or aggregates are found in the 3-20 μm size-fraction (27). In most cases, they are very difficult to culture. Unicellular eukaryotes, or protists, cover a broad range of cell size (0.8-2000 μm). They are taxonomically very diverse with representatives in all of the 8 super-groups of the eukaryotic tree of life (28), whose roles in marine and Earth systems ecology are largely unexplored. Only the most abundant groups, such as diatoms and dinoflagellates, have been studied extensively in the field and cultured successfully (29). Meso-zooplankton (metazoans; multicellular eukaryotes) range in size from 50 μm to tens of metres in colonial forms, and play a pivotal role in both the transfer of energy to higher trophic levels such as fish and other large predators, and in the vertical export of particulate matter produced at the surface of the ocean (30). Their life history (e.g., metabolism, development, locomotion, reproduction, feeding) and their body-size are important properties affecting these two processes (31).

Various sampling methods were used to capture the diversity of both the dominant and less abundant organisms described above (see ***Technical Validation Section***). These methods effectively separated organisms into 10 size fractions: <5 μm (or <3 μm), 5-20 μm (or 3-20 μm), <20 μm, 20-180 μm and 180-2000 μm for planktonic viruses, prokaryotes and unicellular eukaryotes, and >50 μm, >200 μm, >300 μm, >500 μm and >680 μm for large planktonic unicellular eukaryotes and metazoans. Whenever possible, replicate sampling was performed to assess plankton natural variability and to ensure long-term storage of samples in view of future re-analysis using new technologies, notably in the fields of high throughput imaging and – omics which are evolving extremely rapidly.

Detailed protocols concerning the filtration, preservation and storage of plankton samples will be described in detail as a separate publication. Morphological data will be openly released at PANGAEA (<u>http://www.pangaea.de</u>) and nucleotides data will be openly released as they become available at the European Nucleotide Archive (<u>http://www.ebi.ac.uk/ena/</u>).

### [6a] Sampling planktonic viruses, prokaryotes and unicellular eukaryotes

Sampling devices used to collect small size organisms (<20 μm size fractions) include Niskin bottles [NISKIN] mounted on the rosette [RVSS] or occasionally attached individually on the oceanographic cable, a High Volume Peristaltic Pump [HVP-PUMP], and exceptionally a 10-L plastic bucket [BUCKET] (Table 1). Waste waters from *Tara* were not released automatically at sea and were not purged during stations. The pump [HVP-PUMP] was fixed on deck and connected to a 40 mm diameter flexible tube, lowered from either starboard to sample surface water (3-7m) or from the stern to sample other depths down to 60 m. At the beginning of each sampling activity, the tube was rinsed by letting seawater flow overboard for 10 minutes. A single deployment of the pump [HVP-PUMP] brought back up to 1000 L, whereas 4-6 deployments of the rosette [RVSS] [NISKIN] were necessary to collect 400-600 L of seawater.

The choice of the sampling device was determined by weather conditions and the depth of the targeted environmental features. The ***surface water layer*** was systematically sampled using the pump [HVP-PUMP], and exceptionally (at 3 stations) using a 10-Litres plastic bucket [BUCKET]. The ***deep chlorophyll maximum layer*** was sampled preferentially with the pump [HVP-PUMP] or alternatively using multiple deployments of the rosette [RVSS] [NISKIN] when the sampling depth was >60 m. A WETLabs three-optical-sensor [ECOTriplet], measuring pressure, temperature, CDOM and chlorophyll fluorescence, was attached to the intake of the pump tube to monitor sampling depth, temperature and chlorophyll concentrations in real time during pumping. The ***mesopelagic zone*** was systematically sampled using multiple deployments of the rosette [RVSS] [NISKIN].

***Whole seawater*** collected by these devices was then pre-filtered successively on nylon conical sieves with a mesh of 200 μm and 20 μm, and additionally 5 μm for protists (Table 1). The filtrate was collected in four to six 100-L polyethylene containers, which were thoroughly washed with 0.1% bleach, rinsed twice with fresh water and rinsed again twice with the filtrate. Depending on protocols, the <5 μm and <20 μm filtrates were further fractionated on-board using one or a combination of membranes with pore sizes 0.1 μm, 0.2 μm, 0.45 μm, 0.7 μm, 0.8 μm, 1.6 μm or 3 μm. The retention efficiency of meshes, pore-membranes and fibre-filters is a constant debate in plankton ecology. Organisms display various shapes, including high length-to-width ratios, some may easily “squeeze” through pores smaller than their “normal” size, and others may form colonies or tend to aggregate into particles much larger than their individual size. We do not intend to assess the efficiency of the various meshes and filters in retaining the different groups of organisms targeted during the Tara Oceans Expedition. We simply picked commonly used size-fractions and accept the fact that organisms or parts of organisms from the different groups may be present in several size-fractions.

The choice of size thresholds used to collect small eukaryotes, and prokaryotes associated with small particles or with eukaryotes varied during the *Tara* Oceans Expedition, between 3-20 μm and 5-20 μm. Plankton from that size fraction comprise organisms that are often not abundant enough in ***whole seawater*** and often too fragile to be collected with plankton nets that are themselves too delicate to be deployed in rough seas. The sampling method was therefore weather-dependent and often a combination of using either the pump [HVP-PUMP] or Niskin bottles [RVSS] [NISKIN] for ***whole seawater***, and using a plankton net with a mesh size of 5 μm [NET-SINGLE-5] when sea conditions allowed.

When the pump [HVP-PUMP] or Niskin bottles [RVSS] [NISKIN] were used to collect the 3-20 μm or 5-20 μm size-fraction, ***whole seawater*** was pre-filtered successively on 200 μm and 20 μm, and either collected on 3 μm membrane filters (1 to 5 replicates) or concentrated in a conical, 5 μm mesh nylon sieve [SIEVE-5]. The volume of water passing through each 3 μm pore size membrane was consistently 100 L, whereas the volume passing through the [SIEVE-5] was estimated by recording the pumping rate and the start and end time of pumping. Material collected in the [SIEVE-5] was rinsed into an 8-L polyethylene bottle using seawater pre-filtered on 0.1 μm, up to a final volume of 3 L. The 3-20 μm method was used throughout the *Tara* Oceans Expedition (2009-2013), whereas the 5-20 μm method was used only in 2009-2012, i.e. not in the Arctic.

When nets [NET-SINGLE-5] were used for the 5-20 μm size-fraction, they were lowered to the selected environmental feature and towed horizontally for 5-15 min at a speed of 0.3 m/s. Net samples from the cod-end were sieved through 20 μm [SIEVE-20] and poured into an 8-L polyethylene bottle, which was thoroughly pre-washed with 0.1% bleach, rinsed twice with fresh water and rinsed again twice with seawater pre-filtered on 0.1 μm. The volume of ***net sample*** was adjusted to 3 L with 0.1 μm pre-filtered seawater. After each use, nets, cod-ends, and sieves were rinsed with fresh water and checked for holes. That method was used only in 2009-2012, i.e. not in the Arctic.

Plankton from the 20-180 μm size-fraction were collected using a double plankton net with a 20 μm mesh size [NET-DOUBLE-20]. Nets were lowered to the selected environmental feature and towed horizontally for 5-15 min at a speed of 0.3 m/s. Net samples from the two cod-ends were sieved through 180 μm [SIEVE-180] and poured into an 8-L polyethylene bottle, which was thoroughly pre-washed with 0.1% bleach, rinsed twice with fresh water and rinsed again twice with seawater pre-filtered on 0.1 μm. The volume of ***net sample*** was adjusted to 3 L with 0.1 μm pre-filtered seawater. After each use, nets, cod-ends, and sieves were rinsed with fresh water and checked for holes.

Plankton from the 180-2000 μm size-fraction were collected using a 180 μm Bongo Net [NET-BONGO-180]. Nets were lowered to the selected environmental feature and towed horizontally for 5-15 min at a speed of 0.3 m/s. Net samples from the two cod-ends were sieved through 2000 μm [SIEVE-2000] and poured into an 8-L polyethylene bottle, which was thoroughly pre-washed with 0.1% bleach, rinsed twice with freshwater and rinsed again twice with seawater pre-filtered on 0.1 μm. The volume of ***net sample*** was adjusted to 3 L with 0.1 μm pre-filtered seawater. After each use, nets, cod-ends, and sieves were rinsed with fresh water and checked for holes.

### [6b] Sampling large planktonic unicellular eukaryotes and metazoans

Sampling devices used to concentrate and collect the larger and less abundant organisms (>50 μm size fractions) consisted of plankton nets with mesh sizes ranging from 50 to 680 μm [NET-*TYPE-MESH*] and metal pan-shaped sieves [SIEVE-*MESH*] to remove large organisms as needed (Table 1). All nets were equipped with a flow meter and a temperature-depth recorder, and their depth was monitored and adjusted during deployments using an acoustic SCANMAR system. Upon recovery, all nets were rinsed from the outside with running seawater. Cod-ends and metal sieves used to size-fractionate samples were rinsed with running seawater pre-filtered successively on 25 μm and 0.1 μm, using Polygard-CR Cartridge Filters (CR2501006, CRK101006). After each use, nets, cod-ends, and sieves were rinsed with fresh water and checked for holes.

Organism-selectivity and capture-efficiency of plankton nets depend on the mesh size and deployment methods, i.e. depth, tow method (oblique/vertical/horizontal), tow speed and time of day (32). During the *Tara* Oceans Expedition, plankton nets were deployed during day and night in order to capture the nycthemeral vertical migrations. Both [NET-WPII-50] and [NET-WPII-200] were towed vertically or obliquely from a depth of 100 m to the surface, during night and daytime, at a speed of 0.3-0.5 m/s depending on weather conditions. Both [NET-BONGO-300] and [NET-REGENT-680] were towed obliquely from a depth of 500 m to the surface, during night and daytime, at a speed of 0.5 m/s depending on weather conditions. Net samples were preserved on-board with buffered formaldehyde, ethanol or RNA-Later for later morphological and/or molecular analyses.

Where time and weather allowed, a multiple opening-closing net equipped with 5 nets of 300 μm mesh size [NET-MULTI-300] was deployed preferentially at night or during daytime to study the vertical distribution of zooplankton. Nets opened and closed at selected depths between 1000 m and the surface, according to water column features identified from vertical profiles of temperature, salinity, fluorescence, nutrients, oxygen, and particulate matter. Samples were preserved on-board in buffered formaldehyde. The Underwater Vision Profiler (UVP) mounted on the rosette was also used to study the vertical distribution of zooplankton >600 μm during day and night.

In 2011-2013, a neuston net [NET-MANTA-500] was towed at the surface for about 1h at a speed of 0.7 m/s in order to collect plastic particles and associated organisms. Samples were preserved in ethanol for later morphological and molecular analyses. Finally, a Continuous Plankton Recorder (CPR) was deployed between stations in 2013. Samples were preserved in formaldehyde and sent to the Sir Alister Hardy Foundation for Ocean Science (SAHFOS) for later morphological and molecular analyses.

## Data Records

*Tara* Oceans developed best practices for the standardisation and interoperability of data generated across environmental, morphological and molecular analyses. This effort contributed to the publication of a set of standards for reporting and serving data in Marine Microbial Biodiversity, Bioinformatics and Biotechnology (M2B3) (33). Here we describe three levels of the M2B3 reporting standard: ***campaigns***, ***stations*** and ***events***. For each level, we provide a registry of all campaigns/stations/events, pdf documents describing each campaign/station/event, and universal resource locator (URL) queries to access related nucleotides and environmental data.

### Labels

Labels that identify each campaign, station or event are built using a consistent syntax that has the following format:

***Campaign*** labels: TARA_date(yyyymmddZ), e.g.,TARA_20110401Z

***Station*** labels: TARA_station#(001-210), e.g.,TARA_100

***Event*** labels: TARA_datetime(yyyymmddThhmmZ)_station#(001-210)_EVENT_TYPE, e.g.,TARA_20110416T1306Z_100_EVENT_CAST

### Registries

The Tara Oceans Expedition (2009-2013) comprised 60 campaigns (from port-to-port), 210 stations and over 3200 sampling events. Three registries listing the campaigns, stations and events are published at PANGAEA, Data Publisher for Earth and Environmental Science:

***Campaigns*** registry (Data Citation 6): <u>http://doi.pangaea.de/10.1594/PANGAEA.842191</u>

***Stations*** registry (Data Citation 7): <u>http://doi.pangaea.de/10.1594/PANGAEA.842237</u>

***Events*** registry (Data Citation 8): <u>http://doi.pangaea.de/10.1594/PANGAEA.842227</u>

The ***campaigns*** registry provides details about the scientific interest of each campaign, a list of scientists on board, and URLs for the corresponding campaign summary report (pdf), environmental data sets and nucleotides data sets.

The ***stations*** registry provides details about the geographic context of each station, including mean and maximum bathymetric depth (extracted from the General Bathymetric Chart of the Oceans; GEBCO), minimum distance from the coast, the corresponding marine biomes and biogeographical provinces defined by Longhurst (34), and when applicable information about the corresponding exclusive economic zone and related legal aspects. Additionally for each station, we provide information about the environmental features that were sampled and their depth, and the number of deployments carried out with the different sampling devices listed in Table 1. URLs provide access to the corresponding oceanographic context report (pdf), environmental data sets and nucleotides data sets.

The ***events*** registry provides details about the sampling date, time, location and methodology of each event. URLs provide access to the corresponding event log sheet (pdf), environmental data sets and nucleotides data sets. Sampling events occurring outside the context of a station were assigned the station label TARA_999. Such events include for example underway measurements of the on-board meteorological station [BATOS] and of the continuous surface sampling system [CSSS], and exceptional deployments of plankton nets [NET-*TYPE-MESH*] or of the rosette vertical sampling system [RVSS].

All available metadata information about a single campaign/station/event can be extracted from the respective registries using a web service (description: <u>https://ws.pangaea.de/dds-fdp/</u>). Here we provide an example for each registry:

***Campaign***-specific metadata query: https://ws.pangaea.de/dds-fdp/rest/panquery?datasetDOI=doi.pangaea.de/10.1594/PANGAEA.842191&filterParameter=Campaign&filterValue=TARA_20110401Z

***Station***-specific metadata query: <u>https://ws.pangaea.de/dds-fdp/rest/panquery?datasetDOI=doi.pangaea.de/10.1594/PANGAEA.842237&filterParameter=Station&filterValue=TARA_100</u>

***Event-***specific metadata query: https://ws.pangaea.de/dds-fdp/rest/panquery?datasetDOI=doi.pangaea.de/10.1594/PANGAEA.842227&filterParameter=Event&filterValue=TARA_20110416T1306Z_100_EVENT_CAST

Additionally, the same web service can be used to extract information about all stations from a campaign, all events from a campaign, or all events from a station as shown in these examples:

***Stations-from-campaign***-specific metadata query: https://ws.pangaea.de/dds-fdp/rest/panquery?datasetDOI=doi.pangaea.de/10.1594/PANGAEA.842237&filterParameter=Campaign&filterValue=TARA_20110401Z

***Events-from-a-campaign***-specific metadata query: https://ws.pangaea.de/dds-fdp/rest/panquery?datasetDOI=doi.pangaea.de/10.1594/PANGAEA.842227&filterParameter=Campaign&filterValue=TARA_20110401Z

***Events-from-a-station***-specific metadata query: https://ws.pangaea.de/dds-fdp/rest/panquery?datasetDOI=doi.pangaea.de/10.1594/PANGAEA.842227&filterParameter=Station&filterValue=TARA_100

### Reports and log sheets

Campaign reports were written by the chief scientist and scientific crew in order to document the objectives and main achievements of each campaign, as well as any deviations from the regular sampling programme (e.g.,<u>TARA_20110401Z_report.pdf</u>). The station reports were written by Flavian Kokoszka, Rémi Laxenaire and Sabrina Speich to provide background knowledge of the physical oceanography at each station (e.g.,<u>TARA_100_oceanographic_context_report.pdf</u>). Finally, event log sheets were filled on board each time a sampling device was deployed, recording the position, date, time, type of device, sampling depths, volume sampled, ID and filename of sensor outputs, operator’s comments, and unique identifiers (barcodes) of samples collected during the event (e.g.,<u>TARA_20110415T1312Z_100_EVENT_CAST.pdf</u>). Most of the information found in reports and log sheets was extracted manually, quality checked using controlled vocabularies, and archived in the campaigns, stations and events registries. Nevertheless, these narrative documents remain a valuable and complementary source of information. The three registries contain universal resource locators (URLs) pointing to the reports and log sheets of each campaign/station/event. Reports and log sheets can also be browsed directly in the PANGAEA store:

***Campaign*** reports: <u>http://store.pangaea.de/Projects/TARA-OCEANS/Campaign_Reports/</u>

***Station*** reports: <u>http://store.pangaea.de/Projects/TARA-OCEANS/Station_Reports/</u>

***Event*** log sheets: <u>http://store.pangaea.de/Projects/TARA-OCEANS/Logsheets_Event/</u>

### Up-to-date lists of available nucleotides and environmental data sets

Tara Oceans data will progressively be released openly following their analysis and validation. Up-to-date lists of nucleotides and environmental data sets can be obtained from universal resource locator (URL) queries that are made specific to any campaign/station/event by using labels from the campaigns, stations and events registries.

A list of environmental data sets published at PANGAEA can be obtained by combining the following base URL: <u>http://www.pangaea.de/search?q=</u> with a search term. The URL query is made specific to any Tara Oceans campaign, station or event by adding the corresponding label as the search term, see examples below. Already-built URL queries for each campaign, station or event are provided in the three respective registries (Data Citations 6-8).

***Campaign***-specific environmental data query: <u>http://www.pangaea.de/search?q=TARA_20110401Z</u>

***Station***-specific environmental data query: <u>http://www.pangaea.de/search?q=TARA_100</u>

***Event*** specific environmental data query: <u>http://www.pangaea.de/search?q=TARA_20110416T1306Z_100_EVENT_CAST</u>

A list of nucleotides data published at ENA can be obtained by combining the following base URL: http://www.ebi.ac.uk/ena/data/search?query= with a search term. The URL query is made specific to any Tara Oceans campaign, station or event by adding the corresponding label as the search term, see examples below. Already-built URL queries for each campaign, station or event are provided in the three respective registries (Data Citations 6-8).

***Campaign*** specific nucleotides data query: <u>http://www.ebi.ac.uk/ena/data/search?query=TARA_20110401Z</u>

***Station*** specific nucleotides data query: <u>http://www.ebi.ac.uk/ena/data/search?query=TARA_100</u>

***Event*** specific nucleotides data query: <u>http://www.ebi.ac.uk/ena/data/search?query=TARA_20110415T1245Z_100_EVENT_PUMP</u>

## Technical Validation

Here we provide a first order validation of the *Tara* Oceans sampling methodology by compiling published values of plankton cell/body size, natural abundance and richness (Table 2). These are compared to the sampling volume and mesh size of the different sampling methods (Figure 5).

**Figure 5.**
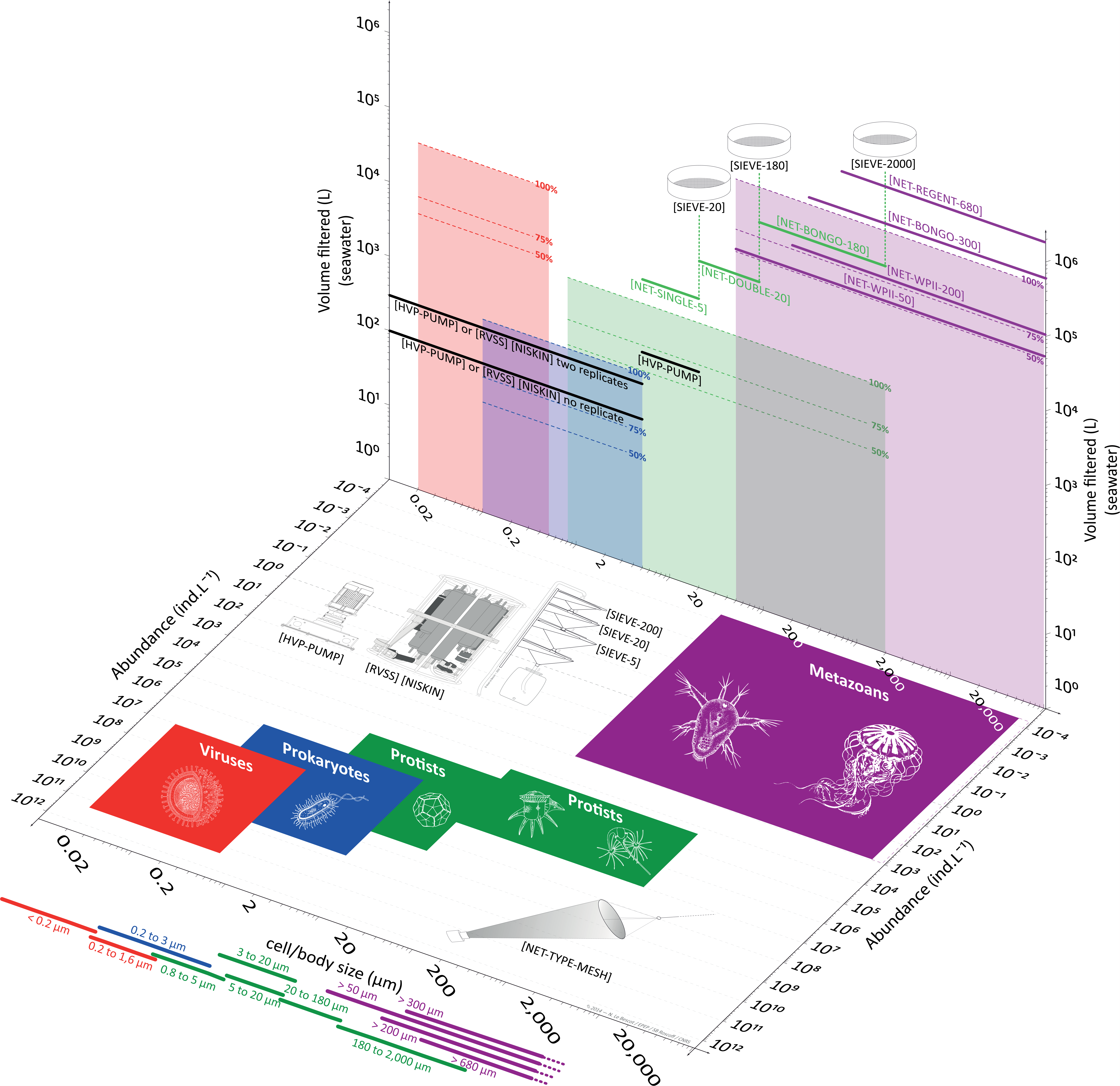
Empirical basis for the size-fractionation approach and the choice of sampling devices. The horizontal plane shows the range of body/cell size and natural abundances reported in the literature (Table 2) for viruses (including giant viruses), prokaryotes, protists and metazoans (coloured boxes). The sampling devices used to collect plankton <5 μm in size (i.e. high volume peristaltic pump and rosette with Niskin bottles) and >5 μm in size (i.e. plankton nets) are illustrated as well on the horizontal plane. The vertical plane shows the volume of seawater required to capture 100%, 75% and 50% of species richness reported in the literature (Table 2) for viruses (including giant viruses), prokaryotes, protists and metazoans (shaded boxes). The typical volume of seawater collected by sampling devices are shown in comparison (horizontal thick lines). Also illustrated on the vertical plane: Sieves were used to remove large organisms in the case of plankton nets for protists (5, 20 and 180 μm mesh).

**Table 2.**
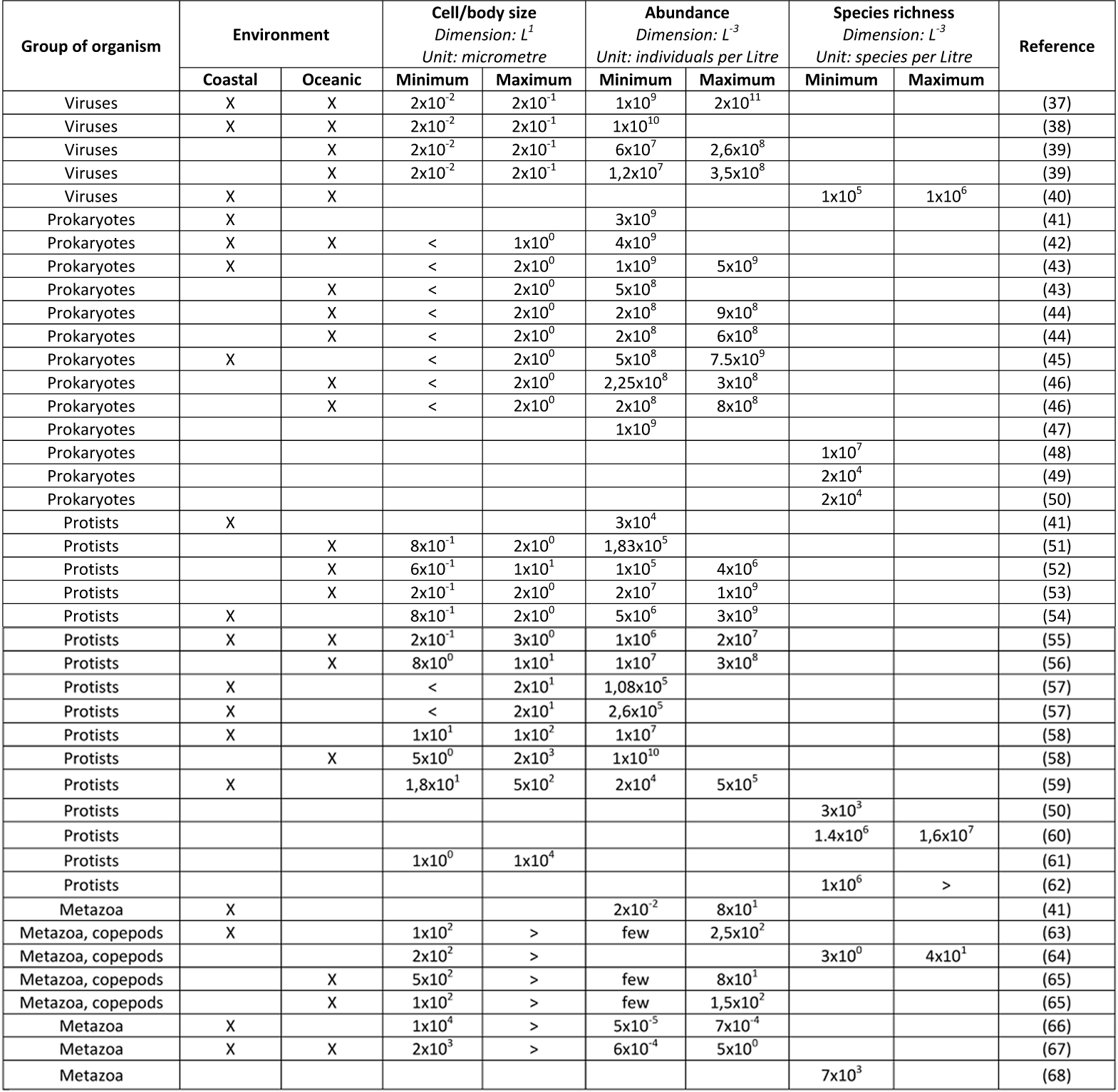
Cell/body size, abundance and species richness reported in the literature for the four groups of plankton in coastal and oceanic environments.

Life history traits such as cell/body size and the natural range of abundance determine the general structure and dynamics of food webs and other ecological networks, across multiple scales of organisation (35) (31). Here we characterise the five groups of plankton by their size and abundance in seawater, using values from the literature (Table 2). The range of these characteristics are summarised for each plankton group using coloured areas on the horizontal plane of Figure 5. As already described for a wide range of organisms (36), the literature shows an inverse relationship between plankton size and abundance in the natural environment, so that small viruses (10^-2^-10° μm) generally form the most abundant group (10^7^-10^11^ ind. L^-1^), whereas the larger metazoans (10^1^-10^5^ μm) are generally the least abundant group (10^-4^-10^3^ ind. L^-1^).

Species richness and evenness are used to estimate species diversity, and should therefore be considered when designing sampling strategies and methodologies for biodiversity studies. Here we characterise the five groups of plankton by their species richness in seawater, using values from the literature (Table 2, Refs. 37-68). From these values we made “back of the envelope” calculations of the volume of seawater required to capture 100%, 75% and 50% of species within each group of plankton (Figure 5; coloured areas on the vertical plane). The effectiveness of our sampling strategy can be assessed by comparing these coloured areas with the sampling volume of the various sampling devices used during the *Tara* Oceans Expedition (horizontal full lines on the vertical plane).

Based on this assessment, it appears that our sampling strategy would capture <50% of total richness for viruses and small size protists (0.8-5 μm). Accordingly, one would need to filter thousands of litres of seawater in order to capture 75% of total richness for these groups. This is both impractical for most field campaigns and dependent on how one defines the currency of richness for these groups, i.e. the concept of species. In all other groups and size fractions, our sampling strategy appears to capture >75% of species richness, and 100% in the case of large size protists and metazoans. It is important to note that data about plankton richness is very scarce in the literature, so that this assessment is only a first approximation. *Tara* Oceans data will undoubtedly contribute to fill this knowledge gap and improve the sampling design of future ocean biodiversity surveys.

## Usage Notes

The *Tara* Oceans data policy follows the open science principle of open access and early release of raw and validated data sets. All data presented here (Data Citations 6-8) are published under the Creative Commons Attribution 3.0 Unported (CC-by 3.0) and must therefore be cited when used in scientific papers, posters or presentations. As with most scholarly publications, data citations have authors, a title, a year of publication and a digital object identifier (see data citations in the Reference Section). Furthermore, we kindly ask to include the ***Tara* Oceans Consortium** in the acknowledgements. When referring to the *Tara* Oceans Data or to the sampling strategy and methodology of the *Tara* Oceans Expedition, please cite the present paper.

## Acknowledgements

We thank the commitment of the following people and sponsors who made this singular expedition possible: CNRS (in particular the Groupement de Recherche GDR3280), European Molecular Biology Laboratory (EMBL), Genoscope/CEA, Fund for Scientific Research – Flanders, VIB, Stazione Zoologica Anton Dohrn, UNIMIB, ANR (projects POSEIDON/ANR-09-BLAN-0348, BIOMARKS/ANR-08-BDVA-003, PROMETHEUS/ANR-09-GENM-031, PROMETHEUS/ANR-09-PCS-GENM-217, TARAGIRUS/ ANR-09-PCS-GENM-218, OCEANOMICS/ANR-11-BTBR-0008, FRANCE GENOMIQUE/ANR-10-INBS-09-08), EU FP7 (MicroB3/No.287589, IHMS/HEALTH-F4-2010-261376, MetaCardis/HEALTH-F4-2012-305312), ERC Advanced Grant Awards to CB (Diatomite: 294823) and PB (CancerBiome: 268985), Spanish Ministry of Science and Innovation grant CGL2011-26848/BOS MicroOcean PANGENOMICS to SGA, JSPS KAKENHI Grant Number 26430184 to HO, FWO, BIO5, Biosphere 2, agne’s b., the Veolia Environment Foundation, Region Bretagne, World Courier, Illumina, Cap L’Orient, the EDF Foundation EDF Diversiterre, FRB, the Prince Albert II de Monaco Foundation, Etienne Bourgois, the *Tara* schooner and its captain and crew. Tara Oceans would not exist without continuous support from 23 institutes (<u>http://oceans.taraexpeditions.org</u>). Special thanks to Cornelia Behrens, Janine Felden, Florentina Münzner, Lucy Schlicht, Adrian Tanara, Sany Tchanra and Marie-Jeanne Pesant for the manual curation of logsheets and archiving data at PANGAEA. We also acknowledge the work of Andree Behnken who developed the dds-fdp web service. All authors approved the final manuscript. This article is contribution number 26 of the *Tara* Oceans Consortium.

The collection of *Tara* Oceans data was made possible by those who contributed to sampling and to logistics during the *Tara* Oceans Expedition: Alain Giese, Alan Deidun, Alban Lazar, Aldine Amiel, Ali Chase, Aline Tribollet, Ameer Abdullah, Amélie Betus, André Abreu, Andres Peyrot, Andrew Baker, Anna Deniaud, Anne Doye, Anne Ghuysen Watrin, Anne Royer, Anne Thompson, Annie McGrother, Antoine Sciandra, Antoine Triller, Aurélie Chambouvet, Baptiste Bernard, Baptiste Regnier, Beatriz Fernandez, Benedetto Barone, Bertrand Manzano, Bianca Silva, Brett Grant, Brigitte Sabard, Bruno Dunckel, Camille Clérissi, Catarina Marcolin, Cédric Guigand, Céline Bachelier, Céline Blanchard, Céline Dimier-Hugueney, Céline Rottier, Chris Bowler, Christian Rouvière, Christian Sardet, Christophe Boutte, Christophe Castagne, Claudie Marec, Claudie Marec, Claudio Stalder, Colomban De Vargas, Cornelia Maier, Cyril Tricot, Dana Sardet, Daniel Bayley, Daniel Cron, Daniele Iudicone, David Mountain, David Obura, David Sauveur, Defne Arslan, Denis Dausse, Denis de La Broise, Diana Ruiz Pino, Didier Zoccola, Édouard Leymarie, Éloïse Fontaine, Émilie Sauvage, Emilie Villar, Emmanuel Boss, Emmanuel G. Reynaud, Éric Béraud, Eric Karsenti, Eric Pelletier, Éric Roettinger, Erica Goetz, Fabien Perault, Fabiola Canard, Fabrice Not, Fabrizio D’Ortenzio, Fabrizio Limena, Floriane Desprez, Franck Prejger, François Aurat, François Noël, Franscisco Cornejo, Gabriel Gorsky, Gabriele Procaccini, Gabriella Gilkes, Gipsi Lima-Mendez, Grigor Obolensky, Guillaume Bracq, Guillem Salazar, Halldor Stefansson, Hélène Santener, Hervé Bourmaud, Hervé Le Goff, Hiroyuki Ogata, Hubert Gautier, Hugo Sarmento, Ian Probert, Isabel Ferrera, Isabelle Taupier-Letage, Jan Wengers, Jarred Swalwell, Javier del Campo, Jean-Baptiste Romagnan, Jean-Claude Gascard, Jean-Jacques Kerdraon, Jean-Louis Jamet, Jean-Michel Grisoni, Jennifer Gillette, Jérémie Capoulade, Jérôme Bastion, Jérôme Teigné, Joannie Ferland, Johan Decelle, Judith Prihoda, Julie Poulain, Julien Daniel, Julien Girardot, Juliette Chatelin, Lars Stemmann, Laurence Garczarek, Laurent Beguery, Lee Karp-Boss, Leila Tirichine, Linda Mollestan, Lionel Bigot, Loïc Vallette, Lucie Bittner, Lucie Subirana, Luis Gutiérrez, Lydiane Mattio, Magali Puiseux, Marc Domingos, Marc Picheral, Marc Wessner, Marcela Cornejo, Margaux Carmichael, Marion Lauters, Martin Hertau, Martina Sailerova, Mathilde Ménard, Matthieu Labaste, Matthieu Oriot, Matthieu Bretaud, Mattias Ormestad, Maya Dolan, Melissa Duhaime, Michael Pitiot, Mike Lunn, Mike Sieracki, Montse Coll, Myriam Thomas, Nadine Lebois, Nicole Poulton, Nigel Grimsley, Noan Le Bescot, Oleg Simakov, Olivier Broutin, Olivier Desprez, Olivier Jaillon, Olivier Marien, Olivier Poirot, Olivier Quesnel, Pamela Labbe-Ibanez, Pascal Hingamp, Pascal Morin, Pascale Joannot, Patrick Chang, Patrick Wincker, Paul Muir, Philippe Clais, Philippe Koubbi, Pierre Testor, Rachel Moreau, Raphaël Morard, Roland Heilig, Romain Troublé, Roxana Di Mauro, Roxanne Boonstra, Ruby Pillay, Sabrina Speich, Sacha Bollet, Samuel Audrain, Sandra Da Costa, Sarah Searson, Sasha Tozzi, Sébastien Colin, Sergey Pisarev, Shirley Falcone, Sibylle Le Barrois d’Orgeval, Silvia G. Acinas, Simon Morisset, Sophie Marinesque, Sophie Nicaud, Stefanie Kandels-Lewis, Stéphane Audic, Stephane Pesant, Stéphanie Reynaud, Thierry Mansir, Thomas Lefort, Uros Krzic, Valérian Morzadec, Vincent Hilaire, Vincent Le Pennec, Vincent Taillandier, Xavier Bailly, Xavier Bougeard, Xavier Durrieu de Madron, Yann Chavance, Yann Depays, Yohann Mucherie.

## Author Contributions

Contributed to writing this paper: Stéphane Pesant, Fabrice Not, Marc Picheral, Stefanie Kandels-Lewis and Noan Le Bescot. Contributed to sampling planning during the Tara Oceans Expedition (2009-2013): Gabriel Gorsky, Daniele Iudicone, Eric Karsenti, Stefanie Kandels-Lewis, Fabrice Not, Stéphane Pesant, Sabrina Speich, Romain Troublé. Céline Dimier, Marc Picheral, and Sarah Searson contributed extensively to sampling during the *Tara* Oceans Expedition and refined the sampling methods on board. Stefanie Kandels-Lewis contributed as coordinator of scientific operations and logistics. Stéphane Pesant contributed as coordinator of data management. Eric Karsenti contributed as scientific director of the Tara Oceans Consortium. Tara Oceans Coordinators contributed intellectually to this work.

### *Tara* Oceans Coordinators

Silvia G. Acinas^15^, Peer Bork^7^, Emmanuel Boss^16^, Chris Bowler^10^, Colomban De Vargas^3,4^, Michael Follows^17^, Gabriel Gorsky^5,6^, Nigel Grimsley^18,19^, Pascal Hingamp^20^, Daniele Iudicone^9^, Olivier Jaillon^21,22,23^, Stefanie Kandels-Lewis^7,8^, Lee Karp-Boss^16^, Eric Karsenti^8,10^, Uros Krzic^24^, Fabrice Not^3,4^, Hiroyuki Ogata^25^, Stephane Pesant^1,2^, Jeroen Raes^26,27,28^, Emmanuel G. Reynaud^29^, Christian Sardet^30^, Mike Sieracki^31^, Sabrina Speich^11,12^, Lars Stemmann^5^, Matthew B. Sullivan^32^, Shinichi Sunagawa^7^, Didier Velayoudon^33^, Jean Weissenbach^21,22,23^, Patrick Wincker^21,22,23^.

^15^ Department of Marine Biology and Oceanography, Institute of Marine Science (ICM)-CSIC, 08003, Barcelona, Spain.

^16^ School of Marine Sciences, University of Maine, Orono, ME 04469, USA.

^17^ Department of Earth, Atmospheric and Planetary Sciences, Massachusetts Institute of Technology, Cambridge, MA 02139, USA.

^18^ CNRS UMR 7232, Biologie Intégrative des Organismes Marins (BIOM), 66650, Banyuls-sur-Mer, France.

^19^ Sorbonne Universités, Observatoire Océanologique de Banyuls-sur-Mer (OOB), UPMC Paris 06, 66650, Banyuls-sur-Mer, France.

^20^ Aix Marseille Université, CNRS, IGS UMR 7256, 13288 Marseille Cedex 09, France.

^21^ CEA, Genoscope, 91000, Evry France.

^22^ CNRS, UMR 8030, 91000, Evry, France.

^23^ Université d’Evry, UMR 8030, 91000, Evry, France.

^24^ Cell Biology and Biophysics, European Molecular Biology Laboratory, 69117, Heidelberg, Germany.

^25^ Institute for Chemical Research, Kyoto University, 611-0011, Kyoto, Japan.

^26^ Department of Microbiology and Immunology, Rega Institute KU Leuven, 3000, Leuven, Belgium.

^27^ VIB Center for the Biology of Disease, VIB, 3000, Leuven, Belgium.

^28^ Laboratory of Microbiology, Vrije Universiteit Brussel, 1050, Brussels, Belgium.

^29^ School of Biology and Environmental Science, University College Dublin, Dublin 4, Ireland.

^30^ CNRS, UMR 7009, BioDev, Observatoire Océanologique de Villefranche-sur-Mer (OOV), 06230, Villefranche/mer, France.

^31^ Bigelow Laboratory for Ocean Sciences, East Boothbay, ME 04544, USA.

^32^ Department of Ecology and Evolutionary Biology, University of Arizona, Tucson, AZ 85719, USA.

^33^ DVIP Consulting, 92310, Sèvres, France.

## Reference

1. Gross, L. Untapped bounty: sampling the seas to survey microbial biodiversity. PLoS Biol. 5, e85 (2007).

2. Laursen, L. Spain’s ship comes in. Nature 475, 16–17 (2011).

3. Karsenti, E. et al. A holistic approach to marine eco-systems biology. PLoS Biol. 9, e1001177 (2011).

4. Hingamp, P. et al. Exploring nucleo-cytoplasmic large DNA viruses in Tara Oceans microbial metagenomes. The ISME Journal 7, 1678–1695 (2013).

5. Boss, E. et al. The characteristics of particulate absorption, scattering and attenuation coefficients in the surface ocean; Contribution of the Tara Oceans expedition. Methods in Oceanography. 7, 52-62 doi:10.1016/j.mio.2013.11.002 (2013).

6. Roullier, F. et al. Particle size distribution and estimated carbon flux across the Arabian Sea oxygen minimum zone. Biogeosciences 11, 4541–4557 (2014).

7. Brum, J. R. et al. Global patterns and ecological drivers of ocean viral communities. Science doi:10.1126/science.1261498 (2015).

8. de Vargas, C. et al. Eukaryotic plankton diversity in the sunlit global ocean. Science doi:10.1126/science.1261605 (2015).

9. Lima-Mendez, G. et al. Top-down determinants of community structure in the global plankton interactome. Science doi:10.1126/science.1262073 (2015).

10. Sunagawa, S. et al. Structure and Function of the Global Ocean Microbiome. Science doi:10.1126/science.1261359 (2015).

11. Villar, E. et al. Environmental characteristics of Agulhas rings affect inter-ocean plankton transport. Science doi:10.1126/science.1261447 (2015).

12. Testor, P. et al. Gliders as a component of future observing systems, in Proceedings of OceanObs’09: Sustained Ocean Observations and Information for Society (Vol. 2) ESA Publication WPP-306 (2010).

13. Xing, X. et al. Combined processing and mutual interpretation of radiometry and fluorimetry from autonomous profiling Bio-Argo floats: Chlorophyll a retrieval. J. Geophys. Res. 116, C06020 (2011).

14. Follows, M. J., Dutkiewicz, S., Grant, S. & Chisholm, S. W. Emergent biogeography of microbial communities in a model ocean Science 315, 1843–1846 (2007).

15. Slade, W. H. et al. Underway and moored methods for improving accuray in measurement of spectral particulate absorption and attenuation. J. Atmos. Oceanic Technol. 27 (10), 1733–1746 (2010).

16. Falkowski, P. G. Physiological responses of phytoplankton to natural light regimes. J. Plankton Res. 6 (2), 295–307 (1984).

17. Buttigieg, P. L. et al. The environment ontology: contextualising biological and biomedical entities. J. Biomed. Semant. 4 (43) (2013).

18. Picheral, M. et al. The Underwater Vision Profiler 5: An advanced instrument for high spatial resolution studies of particle size spectra and zooplankton. Limnology and Oceanography: Methods 8 (9), 462-473 doi:10.4319/lom.2010.8.462 (2010).

19. Arrigo, K. R. Marine microorganisms and global nutrient cycles. Nature 437, 349–355 (2005).

20. Falkowski, P. G., Fenchel, T. & Delong, E. F. The microbial engines that drive Earth’s biogeochemical cycles. Science 320, 1034–1039 (2008).

21. Karl, D. M. Microbial oceanography: paradigms, processes and promise. Nat. Rev. Microbiol. 5, 759–769 (2007).

22. Breitbart, M. Marine viruses: truth or dare. Ann. Rev. Mar. Sci. 4, 425–448 (2012).

23. Thomas, R. et al. Acquisition and maintenance of resistance to viruses in eukaryotic phytoplankton populations. Environmental Microbiology 13, 1412–1420 (2011).

24. Monier, A., Claverie, J. M. & Ogata, H. Taxonomic distribution of large DNA viruses in the sea. Genome Biology 9, R106 (2008).

25. Claverie J.-M. & Ogata, H. Ten good reasons not to exclude giruses from the evolutionary picture. Nat. Rev. Microbiol. 7, 615 (2009).

26. Del Giorgio, P. A. & Duarte, C. M. Respiration in the open ocean. Nature 420, 379–384 (2002).

27. Acinas, S. G., Anton, J. & Rodriguez-Valera, F. Diversity of free-living and attached bacteria in offshore Western Mediterranean waters as depicted by analysis of genes encoding 16S rRNA. Appl. Environ. Microbiol. 65 (2), 514–22 (1999).

28. Baldauf, S. L. The Deep Roots of Eukaryotes. Science 300, 1703-1706 doi:10.1126/science.1085544 (2003).

29. Vaulot, D., Le Gall, F., Marie, D., Guillou, L. & Partensky, F. The Roscoff Culture Collection (RCC): a collection dedicated to marine picoplankton. Nova Hedwigia 79, 49–70 (2004).

30. Banse, K. Reflections about chance in my career, and on the top-down regulated world. Ann. Rev. Mar. Sci. 5, 1–19 (2013).

31. Falkowski, P. Ocean Science: The power of plankton. Nature 483, 17–20 (2012).

32. Wiebe, P. H. & Benfield, M. C. From the Hensen net toward four-dimensional biological oceanography. Progress in Oceanography 56, 7–136 (2003).

33. Ten Hoopen, P. et al. Marine microbial biodiversity, bioinformatics and biotechnology (M2B3) data reporting and service standards. Standards in Genomic Sciences doi:10.1186/s40793-015-0001-5.

34. Longhurst, A. Ecological Geograhy of the Sea. (London, 2007).

35. Woodward, G. et al. Body size in ecological networks. Trends Ecol. Evol. 20, 402–409 (2005).

36. Peters, R. H. The Ecological Implications of Body Size. (Cambridge University Press, 1983).

37. Brussaard, C. P., Payet, J. P., Winter, C. & Weinbauer, M. G. Quantification of aquatic viruses by flow cytometry. Manual of Aquatic Viral Ecology 11, 102-107 doi:10.4319/mave.2010.978-0-9845591-0-7.102 (2010).

38. Suttle, C. A. Marine viruses — major players in the global ecosystem. Nat. Rev. Microbiol. 5, 801–812 (2007).

39. Weinbauer, M. G., Rowe, J. M. & Wilhelm, S. W. Determining rates of virus production in aquatic systems by the virus reduction approach. Manual of Aquatic Viral Ecology 1, 1-8 doi:10.4319/mave.2010.978-0-9845591-0-7.1 (2010).

40. Angly, F. E. et al. The marine viromes of four oceanic regions. PLoS Biology 4, e368 (2006).

41. Gilabert, J. Short-term variability of the planktonic size structure in a Mediterranean coastal lagoon. J. Plankton Res. 23, 219–226 (2001).

42. Buitenhuis, E. T. et al. Bacterial biomass distribution in the global ocean. Earth System Science Data 4, 37–46 (2012).

43. Ducklow, H. W. Bacterial production and biomass in the oceans. Microbial Ecology of the Oceans 1, 85–120 (2000).

44. Hyun, J.-H. & Kim, K.-H. Bacterial abundance and production during the unique spring phytoplankton bloom in the central Yellow Sea. Mar. Ecol. Prog. Ser. 252, 77–88 (2003).

45. Li, W. K. W. Annual average abundance of heterotrophic bacteria and synechococcus in surface ocean waters. Limnol. Oceanogr. 43, 1746–1753 (2007).

46. Church, M. J. Resource control of bacterial dynamics in the sea. Microbial Ecology of the Oceans 1, 335–382 (2008).

47. Scanlan, D. J. et al. Ecological genomics of marine picocyanobacteria. Microbiol. Mol. Biol. Rev. 73, 249–299 (2009).

48. Pedrós-Alió, C. Marine microbial diversity: can it be determined? Trends in Microbiology 14 (6) (2006).

49. Pedrós-Alió, C. The Rare Bacterial Biosphere. Annu. Rev. Marine. Sci. 4, 449–466 (2012).

50. Amaral-Zettler, L. A. et al. Microbial community structure across the tree of life in the extreme Río Tinto. The ISME Journal 5, 42–50 (2011).

51. Raghukumar, S. Ecology of the marine protists, the Labyrinthulomycetes (Thraustochytrids and Labyrinthulids). European Journal of Protistology 38, 127–145 (2002).

52. Signorini, S. R. & McClain, C. R. Environmental factors controlling the Barents Sea spring-summer phytoplankton blooms. Geophys. Res. Lett. 36, 10604 (2009).

53. Evans, C., Archer, S. D., Jacquet, S. & Wilson, W. H. Direct estimates of the contribution of viral lysis and microzooplankton grazing to the decline of a Micromonas spp. population. Aquat. Microb. Ecol. 30, 207–219 (2003).

54. Countway, P. D. & Caron, D. A. Abundance and distribution of Ostreococcus sp. in the San Pedro Channel, California, as revealed by quantitative PCR. Applied and Environmental Microbiology 72, 2496 (2006).

55. Worden, A. Z & Not, F. Ecology and Diversity of Picoeukaryotes in Microbial Ecology of the Oceans (ed David L. Kirchman) 159–205 (John Wiley & Sons, Inc., Hoboken, NJ, USA, 2008).

56. Holligan, P. M. et al. A biogeochemical study of the coccolithophore, Emiliania huxleyi, in the North Atlantic. Geophys. Res. Lett. 7, 879 (1993).

57. Ya-Hui G. et al. Marine Nanoplanktonic diatoms from the coastal waters of Hong Kong. In Proceedings of an International Workshop Reunion Conference, 21-26, 103–104 (2001).

58. Taylor, F. J. R. & Pahlinger, U. The Biology of Dinoflagellates. Botanical Monographs 21, 399–529 (Wiley-Blackwell, 1987).

59. Smalley, G. W. & Coats, D. W. Ecology of the Red-Tide Dinoflagellate Ceratium furca: distribution, mixotrophy, and grazing impact on ciliate populations of Chesapeake Bay. J. Eukaryot. Microbiol. 49, 63–73 (2002).

60. Adl, S. M. et al. Diversity, nomenclature, and taxonomy of protists. Systematic Biology 56, 684 (2007).

61. Behnke, A. et al. Depicting more accurate pictures of protistan community complexity using pyrosequencing of hypervariable SSU rRNA gene regions. Environmental Microbiology 13(2), 340–349 (2010).

62. Edgcomb, V. et al. Protistan microbial observatory in the Cariaco Basin, Caribbean. I. Pyrosequencing vs Sanger insights into species richness. The ISME Journal 6, 1–13 (2011).

63. Turner, J. T. The importance of small planktonic copepods and their roles in pelagic marine food webs. Zool. Stud. 43, 255–226 (2004).

64. Rombouts, I. et al. Global latitudinal variations in marine copepod diversity and environmental factors. Proceedings of the Royal Society B-Biological Sciences 276 (1670), 3053–3062 (2009).

65. Gallienne, C.P. & Robbins, B. D. Is Oithona the most important copepod in the world’s oceans? J. Plankton Res. 23 (12), 1421–1432 (2001).

66. Stemmann, L. et al. Global zoogeography of fragile macrozooplankton in the upper 100–1000 m inferred from the underwater video profiler. ICES Journal of Marine Science: Journal du Conseil 65, 433–442 (2008).

67. Moriarty, R., Buitenhuis, E. T., Le Quéré, C. & Gosselin, M.-P. Distribution of known macrozooplankton abundance and biomass in the global ocean. Earth System Science Data 5, 241–257 (2013).

68. Bucklin, A., Steinke, D. & Blanco-Bercial, L. DNA Barcoding of Marine Metazoa. Annu. Rev. Marine. Sci. 3, 471-508 (2011)

## Data Citations

1. Météo France, Tara Oceans Consortium, C., Tara Oceans Expedition, P. PANGAEA doi:10.1594/PANGAEA.836312 (2014).

2. Boss, E., et al. PANGAEA doi:10.1594/PANGAEA.836318 (2014).

3. Reverdin, G., Le Goff, H., Tara Oceans Consortium, C. & Tara Oceans Expedition, P.PANGAEA doi:10.1594/PANGAEA.836320 (2014).

4. Picheral, M., et al. PANGAEA doi:10.1594/PANGAEA.836321 (2014).

5. Picheral, M., et al. PANGAEA doi:10.1594/PANGAEA.836319 (2014).

6. Tara Oceans Consortium, C. & Tara Oceans Expedition, P. PANGAEA doi:10.1594/PANGAEA.842191 (2015).

7. Tara Oceans Consortium, C. & Tara Oceans Expedition, P. PANGAEA doi:10.1594/PANGAEA.842237 (2015).

8. Tara Oceans Consortium, C. & Tara Oceans Expedition, P. PANGAEA doi:10.1594/PANGAEA.842227 (2015).

